# Pharmacological characterization of seven human histamine H_3_ receptor isoforms

**DOI:** 10.1101/2023.12.06.570349

**Authors:** Meichun Gao, Mabel E. Dekker, Rob Leurs, Henry F. Vischer

**Affiliations:** Department of Medicinal Chemistry, Amsterdam Institute of Molecular Life Sciences, Faculty of Science, Vrije Universiteit Amsterdam, De Boelelaan 1108, 1081 BV Amsterdam, The Netherlands

**Keywords:** histamine H_3_ receptor, isoforms, constitutive activity, binding selectivity, efficacy, GPCR

## Abstract

The histamine H_3_ receptor (H_3_R) regulates as a presynaptic G protein-coupled receptor the release of histamine and other neurotransmitters in the brain, and is consequently a potential therapeutic target for neuronal disorders. The human H_3_R encodes for seven splice variants that vary in the length of intracellular loop 3 and/or the C-terminal tail but are all able to induce heterotrimeric G_i_ protein signaling. The last two decades H_3_R drug discovery and lead optimization has been exclusively focused on the 445 amino acids-long reference isoform H_3_R-445.

In this study, we pharmacologically characterized for the first time all seven H_3_R isoforms by determining their binding affinities for reference histamine H_3_ receptor agonists and inverse agonists. The H_3_R-453, H_3_R-415, and H_3_R-413 isoforms display similar binding affinities for all ligands as the H_3_R-445. However, increased agonist binding affinities were observed for the three shorter isoforms H_3_R-329, H_3_R-365, and H_3_R-373, whereas inverse agonists such as the approved anti-narcolepsy drug pitolisant (Wakix®) displayed significantly decreased binding affinities for the latter two isoforms. This opposite change in binding affinity of agonist versus inverse agonists on H_3_R-365 and H_3_R-373 is associated with their higher constitutive activity in a cAMP biosensor assay as compared to the other 5 isoforms. The observed differences in pharmacology between longer and shorter H_3_R isoforms should be considered in future drug discovery programs.

## 1. Introduction

Histamine is a neurotransmitter that regulates various physiological processes in the central nervous system (Haas et al., 2008; Panula et al., 2015). Histaminergic neurons are located in the tuberomammillary nucleus of the posterior hypothalamus (Panula et al., 1984; Watanabe et al., 1984). These neurons project widely to different brain areas, interconnecting with other neural systems and resulting in various effects on physiology (Haas et al., 2008; Panula et al., 2015). Consequently, dysregulation of histaminergic function likely contributes to various neurological and psychiatric disorders, including narcolepsy, schizophrenia and Tourette Syndrome (Ercan-Sencicek et al., 2010; Lin et al., 2008; Shan and Swaab, 2022).

Histamine acts through four G protein-coupled receptor (GPCR) subtypes, histamine H_1_, H_2_, H_3_ and H_4_ receptors, each with distinct functions and distribution in the brain (Panula et al., 2015). The histamine H_3_ receptor (H_3_R) was identified in 1983 as an autoreceptor that inhibits histamine release in rat brain (Arrang et al., 1983). The H_3_R also functions as heteroreceptor by inhibiting the release of many other neurotransmitters (Haas et al., 2008; Panula et al., 2015). The recent approval of pitolisant (Wakix®) underscores the therapeutic value of H_3_R in promoting wakefulness in narcoleptic patients (Syed, 2016). Moreover, H_3_R ligands have potential therapeutic applications in a range of neurological and psychiatric conditions (Panula et al., 2015; Provensi et al., 2020; Shan and Swaab, 2022).

The human H_3_R cDNA was cloned in 1999 from a thalamus cDNA library and encodes for a 445 amino acids-long GPCR (H_3_R-445) (Lovenberg et al., 1999). The human H_3_R gene (HRH3; GenBank ID: 11255) is located on chromosome 20 (20q13.33) and consists of four exons separated by three introns (GenBank XM_017027623.2) (Wellendorph et al., 2002) (Fig 1A-B), although intron 3 and exon 4 had originally been annotated as 3’UTR of exon 3 (GenBank NM_007232.3) (Tardivel-Lacombe et al., 2001). Next to H_3_R-445, six additional seven transmembrane (7TM) isoforms have been identified in human and these vary in the length of their intracellular loop (ICL)3 and/or have an 8 amino acid extension at the C-tail due to alternative mRNA splicing (Fig 1C and D) (Coge et al., 2001; Nakamura et al., 2000; Tardivel-Lacombe et al., 2001; Wellendorph et al., 2002). The deletion of 30 (H_3_R-415), 32 (H_3_R-413), 80 (H_3_R-373 and H_3_R-365), and 116 (H_3_R-329) amino acids in ICL3 are thought to result from the retention/deletion of pseudo introns within exon 3 (Tardivel-Lacombe et al., 2001), whereas the 8 amino acid C-terminal extension (H_3_R-453 and H_3_R-373) results from a splice site upstream of the stop codon that splices out 604 bp (i.e. intron 3) (Fig 1C and D) (Wellendorph et al., 2002).

**Fig. 1.**
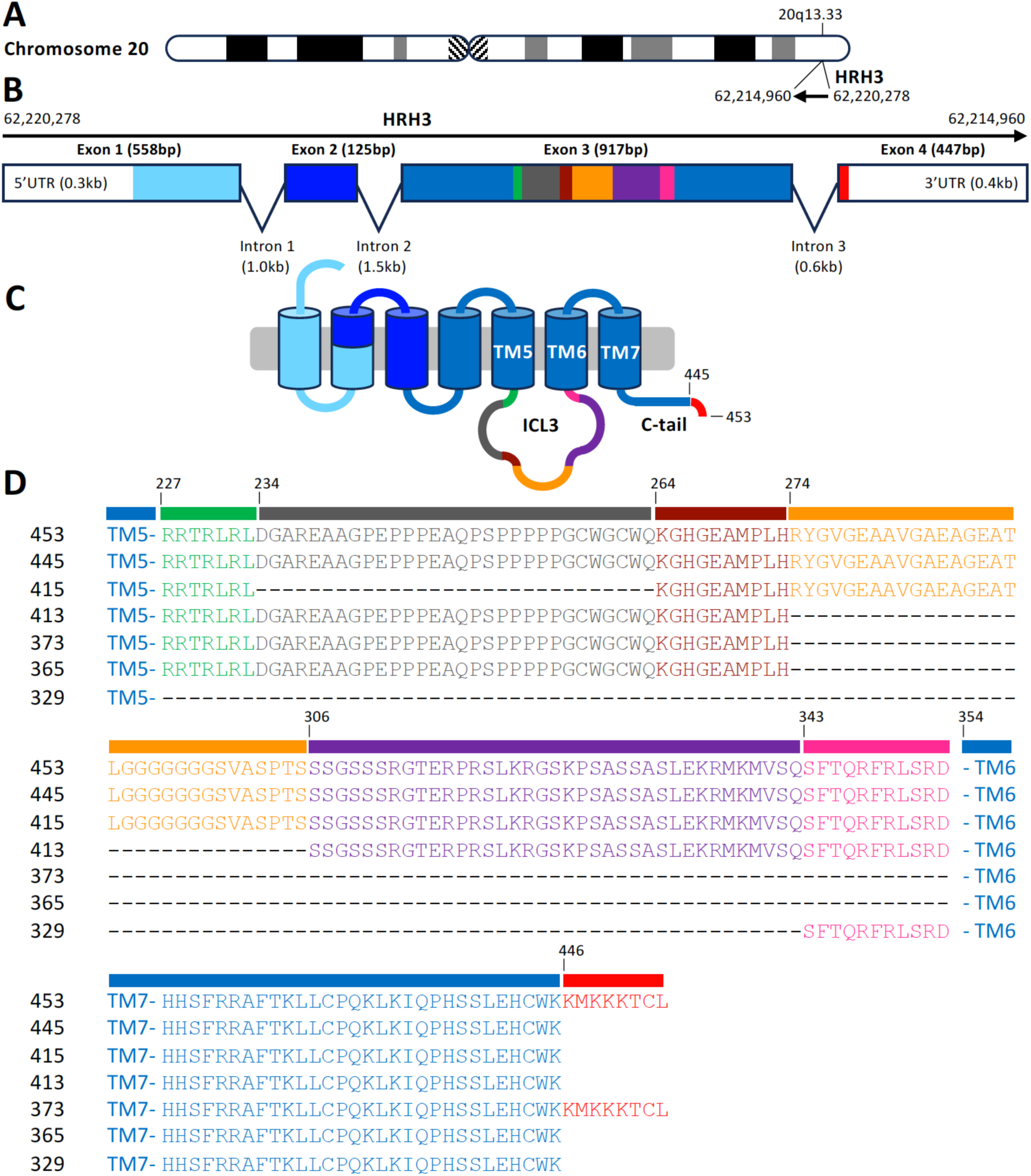
The HRH3 gene and 7TM H_3_R isoforms. (A) Location of HRH3 gene on the reverse strand of chromosome 20 (20q13.33) from nucleotides 62,220,278 to 62,214,960 in the GRCH38.p14 genome assembly (Feb 3, 2022; https://www.ncbi.nlm.nih.gov/datasets/genome/GCF_000001405.40/). (B) HRH3 gene structure (GenBank XM_017027623.2) is depicted by exons in boxes, colors correspond to the H_3_R protein domains that are encoded. (C) Schematic 2D structure of H_3_R-453 protein with the seven transmembrane domain (TM) helices indicated by tubes that are connected by intracellular (ICL) and extracellular loops and an N- and C-terminal tail. The extended C-tail of H_3_R-453 as compared to the canonical H_3_R-445 is indicated in red (D) amino acid sequence alignment of ICL3 between TM5 and TM6, and the C-terminal tail of the seven 7TM H_3_R isoforms with the different segments colored according to the DNA coding sequences. Amino acids numbers are from the H_3_R-445/453 isoforms.

H_3_R drug discovery efforts have been focused on the canonical H_3_R-445 isoform. Except for some basic characterization, the other 7TM H_3_R isoforms have so far been largely understudied. We therefore evaluated the pharmacological properties of 18 H_3_R reference ligands for the first time on all 7TM H_3_R isoforms by determining their binding affinities and efficacies to modulate the canonical H_3_R-mediated inhibition of cAMP production.

## 2. Materials and Methods

### 2.1. Materials

Fetal bovine serum (FBS, #16170078) was obtained from Gibco (Thermo Fisher Scientific). Penicillin/streptomycin (P/S) was purchased from GE Healthcare (Uppsala, Sweden). Dulbecco’s Modified Eagles Medium (DMEM, #41966-029), Dulbecco’s phosphate-buffered saline (DPBS, #15326239), 0.05% Trypsin-EDTA (#11580626), Hanks’ Balanced Salt Solution (HBSS, #11560456) and Pierce^TM^ BCA Protein Assay Kit (#23227) were bought from Thermo Fisher Scientific (Waltham, MA, USA). Linear poly-ethylenimine (PEI, 25-kDa, # 23966-1) was obtained from Polysciences (Warrington, PA, USA). G418 (#108321-42-2) was purchased from Sigma-Aldrich (St. Louis, MO, USA). Zeocin (#ZEL-43-05) was purchase from InvivoGen. Black 96-well plates (#655086) were purchased from Greiner Bio-One (Frickenhausen, Germany). N^α^-[methyl-^3^H] histamine ([^3^H]-NAMH, #NET1027250UC), Microscint-O scintillation liquid (#6013611), GF/C filter plates (#6055690) and MicrobetaWallac Trilux scintillation counter were purchased from PerkinElmer (Groningen, The Netherlands). Histamine (2-(3H-imidazol-4-yl)ethanamine), R-α-methylhistamine (RAMH; (2R)-1-(3H-imidazol-4-yl)propan-2-amine), imbutamine (4-(3H-imidazol-4-yl)butan-1-amine), impentamine (5-(3H-imidazol-4-yl)pentan-1-amine), dimethyl-impentamine (5-(3H-imidazol-4-yl)-N,N-dimethylpentan-1-amine), imetit (2-(3H-imidazol-4-yl)ethylsulfanylmethanimidamide), VUF8328 (3-(1H-imidazol-5-yl)propyl carbamimidothioate), immepip (4-(3H-imidazol-4-ylmethyl)piperidine), methimepip (4-(3H-imidazol-4-ylmethyl)-1-methylpiperidine), immetheridine (4-(3H-imidazol-4-ylmethyl)pyridine), VUF16839 (4-(3-(Propylamino)azetidin-1-yl)pyrimidin-2-amine), pitolisant (1-[3-[3-(4-chlorophenyl)propoxy]propyl]piperidine), clobenpropit (N’-[(4-chlorophenyl)methyl]-1-[3-(3H-imidazol-4-yl)propylsulfanyl]methanimidamide), thioperamide (N-cyclohexyl-4-(3H-imidazol-4-yl)piperidine-1-carbothioamide) and bavisant (JNJ31001074; (4-Cyclopropyl-piperazin-1-yl)-(4-morpholin-4-ylmethyl-phenyl)-methanone) were synthesized in house. Proxyfan (4-[3-(phenylmethoxy)propyl]-3H-imidazole; #2477) was bought from Tocris Bioscience (Bristol, UK). Famotidine (3-[[2-(diaminomethylideneamino)-1,3-thiazol-4-yl]methylsulfanyl]-N’-sulfamoylpropanimidamide; #ab120760) was purchased from Abcam (Cambridge, UK). Mepyramine (N-[(4-methoxyphenyl)methyl]-N’,N’-dimethyl-N-pyridin-2-ylethane-1,2-diamine) was purchased from Research Biochemicals International (Natick, MA, USA). Forskolin (#F-9929) was obtained from LC Laboratories (Woburn, MA, USA). PF-3654746 (N-ethyl-3-fluoro-3-[3-fluoro-4-(pyrrolidin-1-ylmethyl)phenyl]cyclobutane-1-carboxamide; #Axon 1458) and ABT-239 (4-[2-[2-[(2R)-2-methylpyrrolidin-1-yl]ethyl]-1-benzofuran-5-yl]benzonitrile; #Axon 1510) were purchased from Axon Medchem (Groningen, The Netherlands). All these compounds were dissolved in 100% DMSO, except impentamine was dissolved in H_2_O.

### 2.2. Generation of H_3_R-isoform constructs

DNA coding for H_3_R-445 (NM_007232.3) was synthesized by Eurofins with two silent restriction sites (*Eco*RI and *Nhe*I) introduced in TM5 and TM6, respectively, to facilitate the construction of the different isoforms. The H_3_R-445 DNA sequence is preceded by a kozak sequence and flanked by *Kpn*I and *Xba*I restriction sites at its 5’ and 3’ end for subcloning into the mammalian expression vector pcDEF3. DNA fragments coding for the ICL3 sequence of isoforms H_3_R-329, H_3_R-365, H_3_R-413, and H_3_R-415, were synthesized by Eurofins or Biomatik, and subcloned into the H_3_R-445/pcDEF3 plasmid using the silent *Eco*RI and *Nhe*I sites, whereas the synthesized DNA fragment encoding for extended C-terminal tail of H_3_R-373 and H_3_R-453 was subcloned into H_3_R-445/pcDEF3 plasmid using *Nhe*I and *Xba*I. All generated constructs were verified by DNA sequencing.

### 2.3. Cell culture and transfection

HEK293T cells (ATTC, CRL-1573) were cultured in DMEM supplemented with 10% FBS and 1% P/S in a humidified incubator at 37°C with 5% CO_2_. Two million HEK293T cells/dish were seeded one day before transfection with 1 μg H_3_R-isoform plasmid supplemented with 4 μg empty pcDEF3 plasmid using 20 μg linear PEI, as previously described (Mocking et al., 2018). The HEK293 cell line stably expressing a Förster resonance energy transfer (FRET)-based Exchange protein directly activated by cAMP (EPAC) biosensor was kindly provided by Dr. M. Zimmermann (Interax Biotech, Switzerland) (Mathiesen et al., 2013; Scharf et al., 2020). To generate HEK293-EPAC cells stably co-expressing the H_3_R-isoforms, 2 million cells/dish were seeded in 10 cm dishes and transfected the next day with 5 µg H_3_R-isoform plasmid using 20 μg linear PEI. After 2 days, cell culture medium was supplemented with 0.06 mg/ml zeocin and 2 μg/ml G418. Next, monoclonal stable cell lines were generated from 1 cell/well by culturing these cells under these selection conditions.

### 2.4. Radioligand binding assay

Two days after transfection, membranes were prepared from HEK293T cells transiently expressing the H_3_R isoforms. Briefly, cells were collected in ice-cold PBS and centrifuged at 1932 *g* for 10 min at 4°C. Cell pellets were resuspended in Tris-HCl buffer (50 mM, pH 7.4) and disrupted by 5 s sonication. Saturation radioligand binding was performed on membrane suspension with increasing concentration of the radioligand [^3^H]-NAMH for 2 h incubation at 25°C, non-specific binding was determined in the presence of 10 µM clobenpropit. Competition binding was conducted on membrane suspension with 2 nM [^3^H]-NAMH, and increasing concentrations of the tested ligands, for a duration of 2 h at 25°C incubation with continuous shaking at 225 rpm. The incubation process was terminated by harvesting the mixture to 0.5% PEI pre-soaked GF/C filter plates and washed with cold Tris-HCl buffer. After drying for 30 min at 55°C, the radioactivity retained on the filters was quantified using Microbeta Wallac Trilux scintillation counter following the addition of scintillation liquid after a 2 h delay. To determine the receptor expression in EPAC-H_3_R-isoform stable cell lines, cell membranes were directly collected after 24 h growth in 10 cm dish. The maximum number of binding sites (B_max_) were determined by the equilibrium incubation of 2 nM [^3^H]-NAMH in the presence or absence of 10 µM clobenpropit. The protein concentration was subsequently determined by Pierce^TM^ BCA Protein Assay Kit, according to protocol of the manufacturer.

### 2.5. cAMP inhibition by FRET-EPAC biosensor

EPAC-H_3_R-isoform stable cell lines (50,000 cells/well) were seeded into black 96-well plates 1 day before the experiments. The next day, the culture medium was replaced with famotidine (1 μM, diluted in HBSS) and incubated for 30 min at 25°C. Subsequently, 10 µM forskolin was added to enhance basal cAMP levels for an additional 10 min incubation. Fluorescence resonance energy transfer (FRET) measurements were measured in real-time or 90 min after ligand stimulation in a CLARIOstar Plus plate reader (BMG Labtech; Ortenberg, Germany) using excitation at 430-15 nm and emission at 480-20 and 530-20 nm.

### 2.6. Data and statistical analysis

All data are shown as mean ± S.D. of at least three individual experiments. Data and statistical analyses were performed using GraphPad Prism version 9 (GraphPad Software, San Diego, CA, USA). Saturation binding curves were fitted using ‘One-site – Total and nonspecific binding’ to obtain the equilibrium dissociation constant of radioligand (K_d_) and maximum binding sites (B_max_). The equilibrium dissociation constants of unlabeled ligands (K_i_) were calculated from competition binding curves using the Cheng-Prusoff equation by the model of ‘one-site—Fit K_i_’. The receptor expression in EPAC-H_3_R-isoforms stable cell lines, determined from the binding of radioligand at single concentration [A] to receptor, was calculated as below:

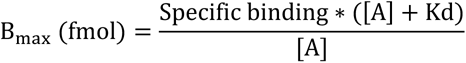

The B_max_ (fmol/mg) was subsequently determined from dividing B_max_ (fmol) by the concentration of protein (mg) from BCA assay.

The signal of cAMP inhibition was quantified by FRET ratio, calculated from dividing the FRET (energy acceptor) signal at 530 nm by the (energy donor) signal at 480 nm. Concentration-response curves were fitted using the model: ‘log(agonist) vs. response (three parameters)’ to obtain EC_50_ and E_max_ values. The response was represented as ΔFRET ratios, obtained by the analysis of ‘Remove baseline and column with the formula ‘Fractional difference: (Value – Baseline)/Baseline’. Intrinsic activities (α) were calculated as maximum response of tested ligand divided by maximum response of reference ligand (i.e. histamine for agonists, and pitolisant for inverse agonists).

## 3. Results

### 3.1. Evaluation of ligand binding affinities for H_3_R isoforms

All seven H_3_R isoforms were transiently expressed in HEK293T-cells and saturation binding of the H_3_R agonist radioligand [^3^H]NAMH was measured on membrane preparations (Supplementary Fig. 1). The radiolabeled agonist bound with high affinity to membranes of HEK293T-cells expressing the various isoforms. The affinity of [^3^H]NAMH for the shorter isoforms H_3_R-373, H_3_R-365 and H_3_R-329 is observed to be higher than for the other isoforms (Table 1).

**Table 1.**
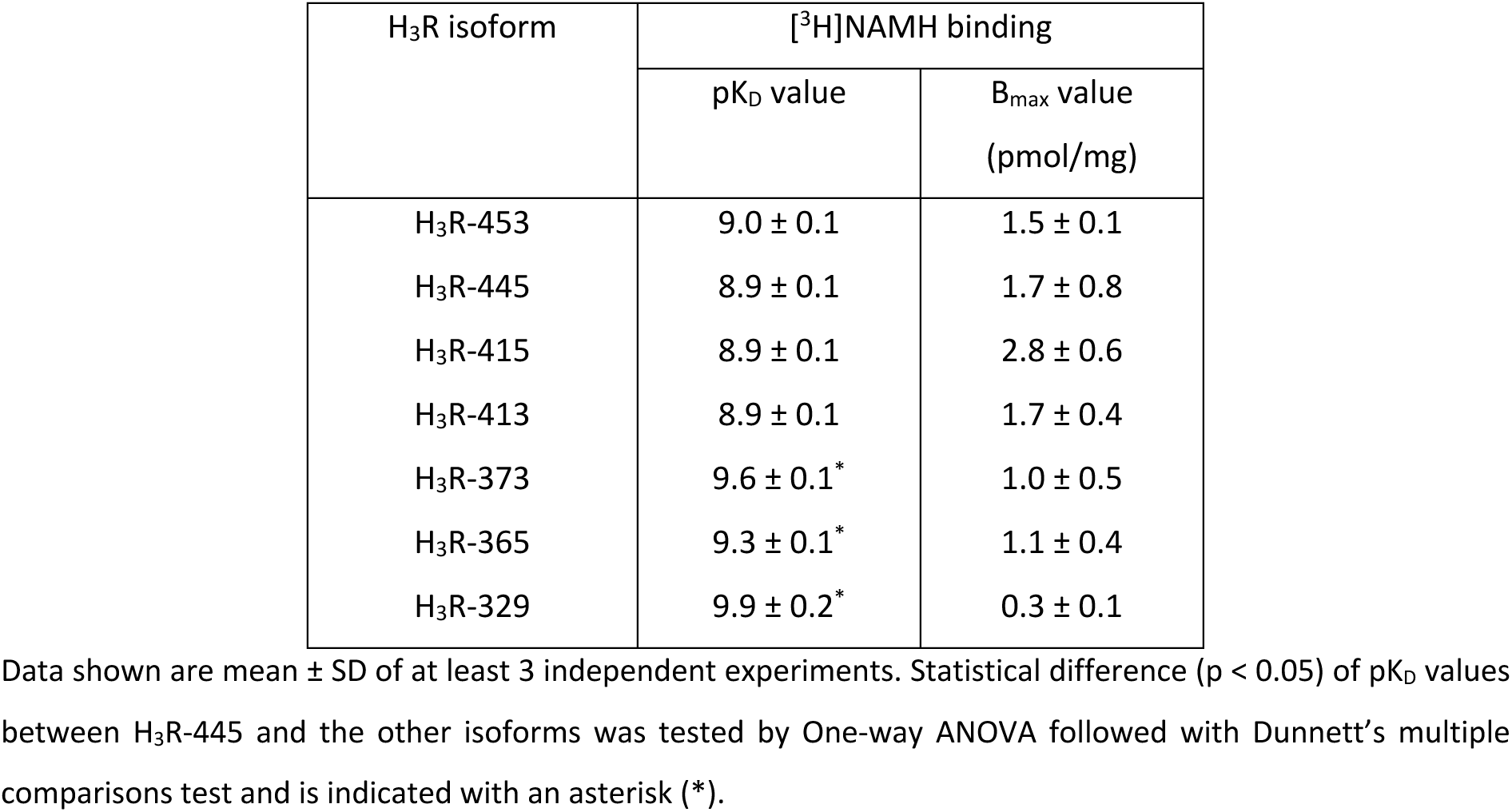
Characterization of [^3^H]NAMH binding to H_3_R isoforms, transiently expressed in HEK293T cells.

Subsequently, the affinity of 18 reference H_3_R agonists and inverse agonists for the seven H_3_R isoforms was measured by radioligand [^3^H]NAMH competition binding assays (Supplementary Fig. 2 and 3). The tested H_3_R agonists display high binding affinities ranging from 0.32 (imetit) to 25 nM (dimethyl-impentamine) for the canonical H_3_R-445 isoform, whereas the natural agonist histamine binds with 8 nM affinity to H_3_R-445 (Table 2, Fig. 2). All tested H_3_R agonists show comparable binding affinities for H_3_R-453, H_3_R-415, and H_3_R-413, as compared to reference isoform H_3_R-445 (Fig. 2A-C). Except for dimethyl-impentamine and impentamine, all other H_3_R agonists showed increased binding affinity for the shorter isoforms H_3_R-373 (Fig. 2D; 3.0- to 6.9-fold) and H_3_R-365 (Fig. 2E; 1.8- to 3.8-fold). Highest binding affinities were observed for all H_3_R agonists on the shortest isoform H_3_R-329 (Fig. 2F; 2.8- to 13.5-fold).

**Fig. 2.**
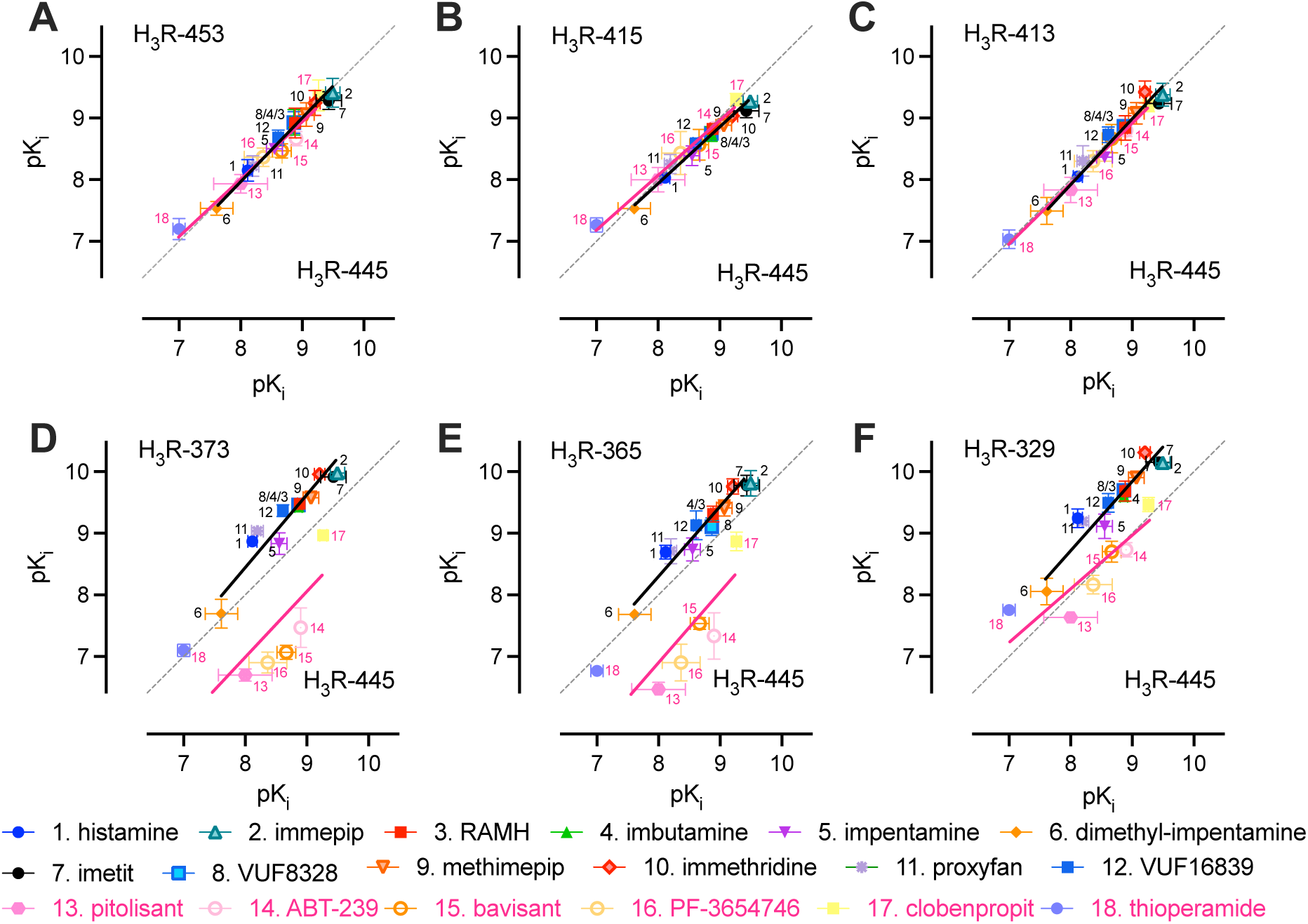
Correlation plots of ligand affinity (pK_i_) values for the H_3_R isoforms. The pK_i_ values were calculated from IC_50_ values extracted from radioligand competition binding curves (sFig 3) using the Cheng-Prusoff correction. The pK_i_ values for the canonical H_3_R-445 isoform are plotted versus pK_i_ values for H_3_R-453 (A), H_3_R-415 (B), H_3_R-413 (C), H_3_R-373 (D), H_3_R-365 (E), and H_3_R-329 (F). H_3_R agonists and inverse agonists/antagonists are indicated by black and magenta numbers, respectively. Deming linear regression was used to compare agonist (black) or inverse agonist/antagonist (magenta) distribution to the (dotted) line of unity. Data are shown as mean ± S.D. of (at least) 3 independent experiments.

**Table 2.**
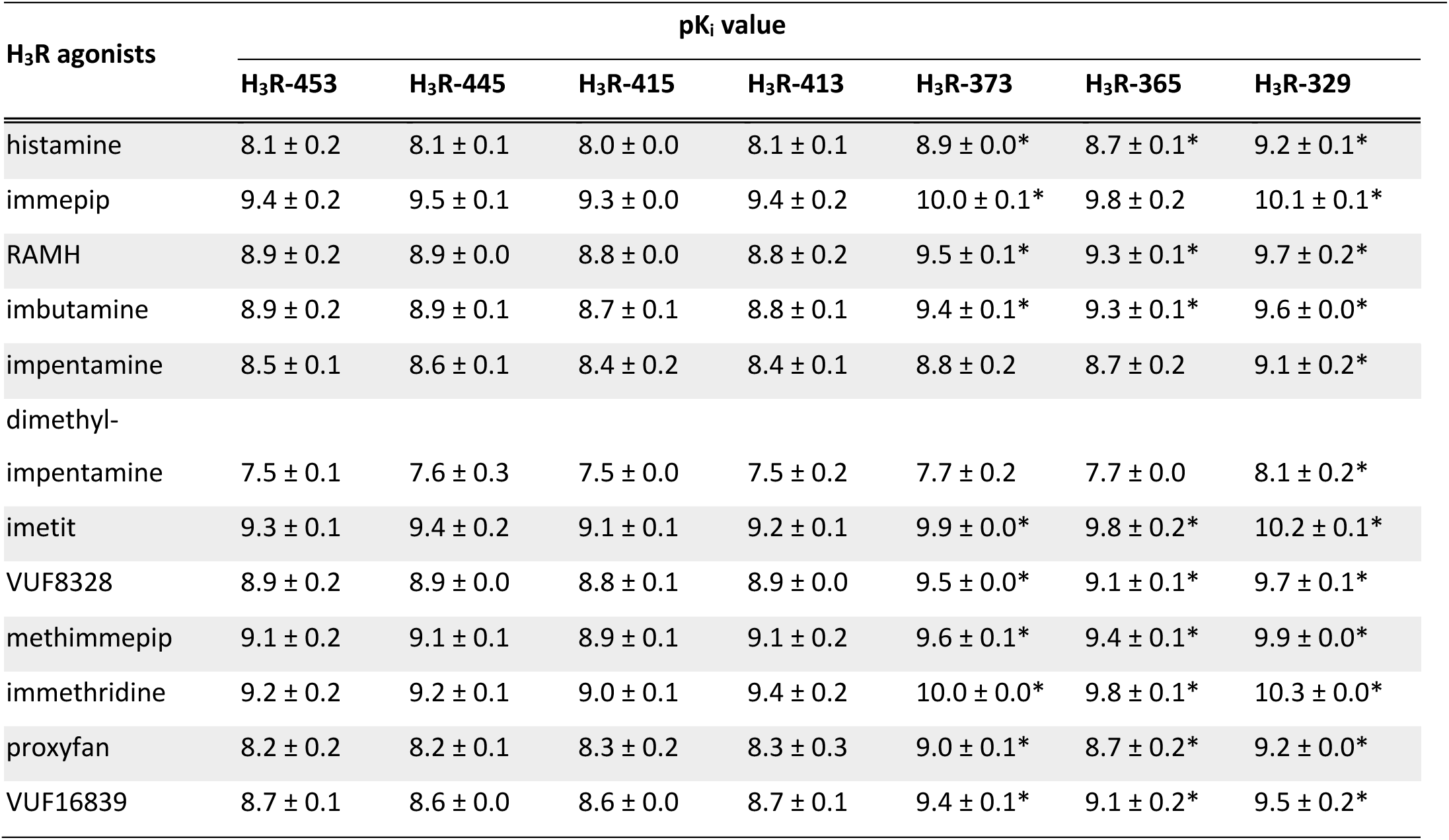
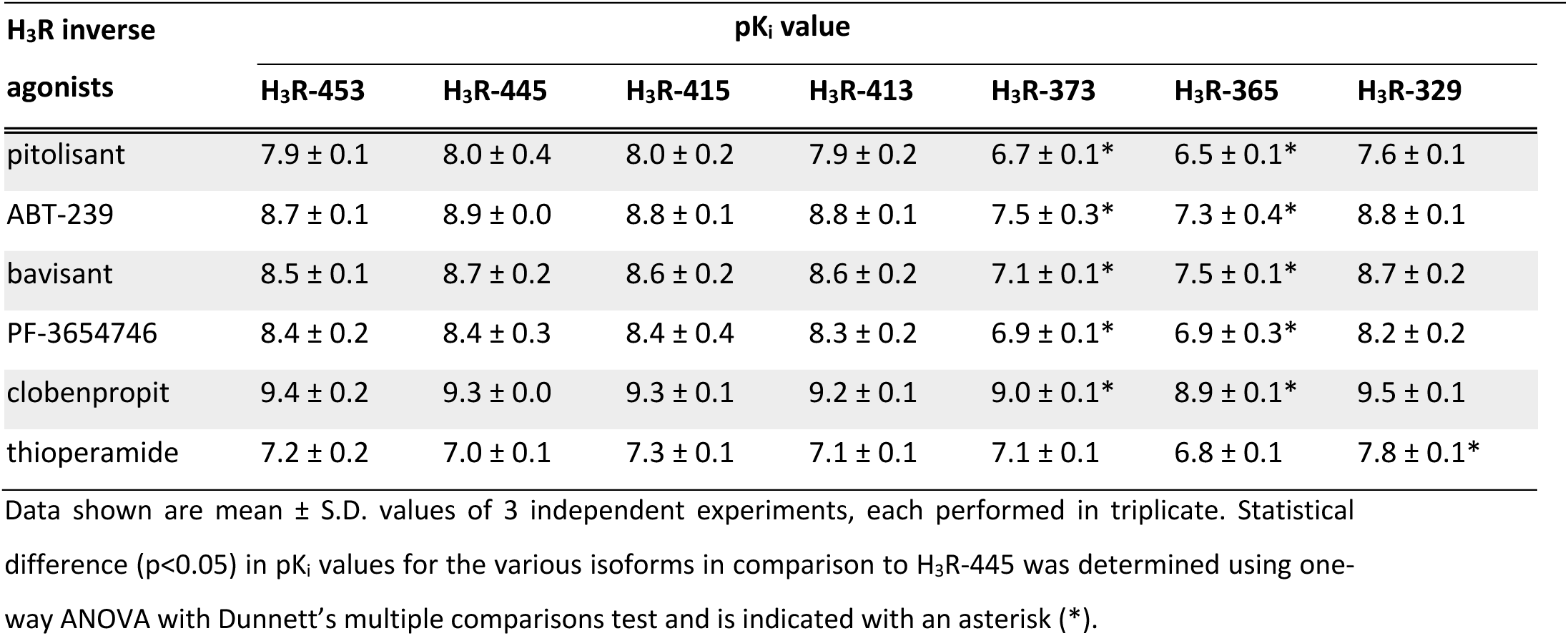
Binding affinities of 18 reference ligands for the H_3_R isoforms expressed in HEK293T cells, by [^3^H]NAMH displacement studies.

The six H_3_R reference inverse agonists/antagonists bound H_3_R-453, H_3_R-415, and H_3_R-413, with comparable affinities as the canonical H_3_R-445 isoform, which is in line with the observations for the 11 tested agonists (Table 2, Fig. 2A-C). In contrast to these agonists, however, pitolisant, ABT-239, bavisant, and PF-3654746 displayed significantly (15- to 40-fold) lower binding affinities for H_3_R-373 and H_3_R-365, whereas the affinity of clobenpropit for both shorter isoforms was only 2- to 2.5-fold lower as compared to H_3_R-445 (Table 2, Fig. 2D-E). Moreover, all tested inverse agonists have similar affinities for the shortest isoform H_3_R-329 as for H_3_R-445, except for thioperamide that showed a 6.8-fold higher affinity for H_3_R-329 in comparison to H_3_R-445 (Table 2, Fig. 2F). In addition, thioperamide bound H_3_R-365 and H_3_R-373 with similar affinity as H_3_R-445 (Table 2, Fig. 2D-E). Hence, the correlation plots show clearly that opposite trends for agonists and inverse agonists (with thioperamide clearly behaving differently).

### 3.2. Inhibition of cAMP production in EPAC biosensor cells stably expressing H_3_R isoforms

The seven H_3_R isoforms were stably expressed in HEK293-EPAC cells to assess their basal and ligand-induced signaling by measuring intracellular cAMP levels using a FRET-based EPAC cAMP biosensor (Scharf et al., 2020). Monoclonal cell lines were generated and selected for H_3_R isoform expression levels of around 500 fmol/mg protein, which can be considered physiologically relevant, being in line with the known expression levels of the H_3_R in e.g. brain tissue (Jansen et al., 2000). Expression levels of most isoforms in these stable cell lines were comparable to H_3_R-445, but H_3_R-453 and H_3_R-365 were expressed at 3.8-fold lower and 1.8-fold higher levels, respectively (Supplementary Table 1).

To measure the canonical inhibition of cAMP production by H_3_R-445 activation upon agonist stimulation, cAMP production was first increased by 10 µM forskolin (FSK), a direct stimulator of adenylate cyclase (Seamon et al., 1981). As expected, histamine initially reduced the FSK-induced cAMP production in a concentration-dependent manner, as detected by an increase in the EPAC biosensor FRET ratio. However, at higher concentrations (> 10 nM) histamine actually increased cAMP production resulting in this biphasic response (Fig. 3A). In contrast, the H_3_R-selective agonist immepip (Vollinga et al., 1994) inhibited the FSK-induced cAMP production in a concentration dependent manner (Fig. 3A), suggesting that histamine might activate endogenously expressed histamine receptor subtypes in the HEK-EPAC cells at higher concentrations and consequently counteracted H_3_R responses. Hence, the HEK-EPAC-445 cells were first pre-incubated with the H_1_R- and/or H_2_R-selective antagonists mepyramine (0.1 µM) and famotidine (1 µM) (Lovenberg et al., 1999; Malone et al., 2021), respectively, before stimulation with 10 µM histamine or immepip (Fig. 3B). Mepyramine did not affect the response of histamine or immepip. On the other hand, famotidine prevented the histamine-induced increase in cAMP levels, without affecting the immepip response, and allowed detection of histamine-induced G_αi_-responses after H_3_R activation in the HEK-EPAC-445. Consequently, famotidine (1 µM) was included in all HEK293-EPAC experiments with the seven H_3_R isoforms to antagonize endogenously expressed H_2_R.

**Fig. 3.**
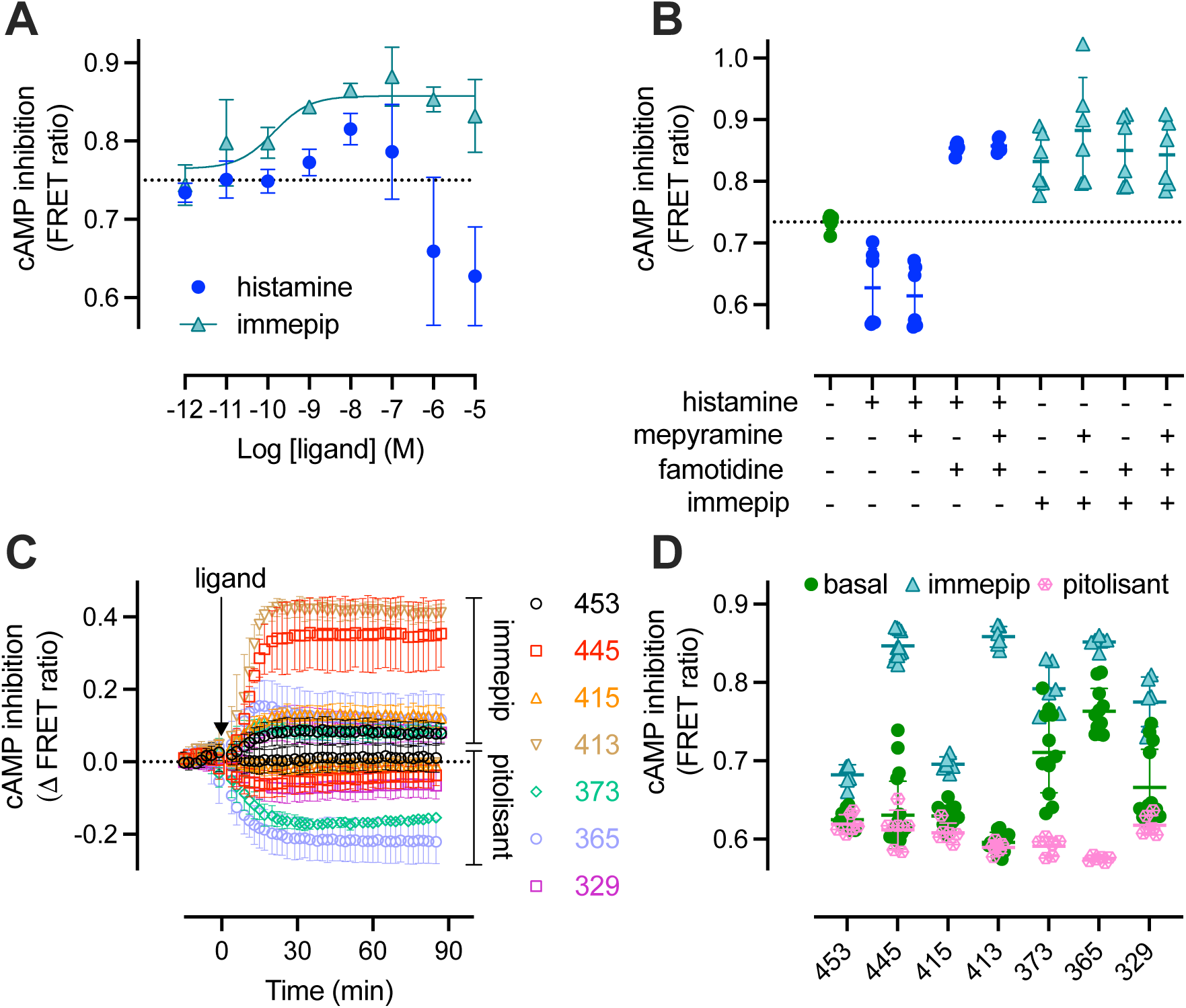
Detection of intracellular cAMP levels by a FRET-based EPAC sensor in HEK293 cells stably co-expressing the H_3_R isoforms. Modulation of forskolin-induced (10 µM) cAMP production in HEK293-EPAC/H_3_R-445 cells by increasing concentrations histamine or immepip (A), or by 10 µM histamine or immepip in combination with 0.1 µM H_1_R-selective antagonist mepyramine and/or 1 µM H_2_R-selective antagonist famotidine (B). Modulation of forskolin-induced (10 µM) cAMP production over baseline (ΔFRET) in HEK293-EPAC co-expressing the various H_3_R isoforms in time following injection of 10 µM agonist immepip or inverse agonist pitolisant in the presence of 1 µM famotidine (C). Detection of basal and ligand-induced (1 µM) modulation of cAMP levels in HEK293-EPAC cells co-expressing the various H_3_R isoforms after 90 min incubation, in the presence of 10 µM forskolin and 1 µM famotidine. Data are the mean ± S.D. of three independent experiments performed. Individual data points are shown in panels B and D with mean indicated by the horizontal line ± S.D.

Next, cAMP levels were measured in real time upon stimulation of the HEK-EPAC cell lines expressing the H_3_R isoforms with the agonist immepip or inverse agonist pitolisant (10 µM). Immepip induced an increase in the ΔFRET ratio of the EPAC biosensor to a steady-state level within approximately 20 min for all H_3_R isoforms, indicating a reduction of cellular cAMP levels as compared to baseline (Fig. 3C). The highest immepip-induced responses were clearly observed for H_3_R-445 and H_3_R-413, whereas the signal was considerably smaller for the other isoforms. Conversely, inverse agonist pitolisant reduced the ΔFRET ratio to a steady-state level within approximately 30 min for H_3_R-373 and H_3_R-365 isoforms and to lesser extent for H_3_R-445, H_3_R-415, and H_3_R-329 (Fig. 3C), suggesting that these isoforms display constitutive signaling in the generated HEK-EPAC cell lines. In contrast, no effect of pitolisant was observed for the cell lines stably expressing H_3_R-413 and H_3_R-453. Indeed, constitutive signaling was clearly observed as increased basal inhibition for H_3_R-373 and H_3_R-365, as compared to H_3_R-445, resulting in a smaller agonist-induced response but a larger inverse agonist-induced response (Fig. 3D). Despite the difference in agonist-induced response windows, comparable cAMP levels (i.e. FRET ratios) were detected in the H_3_R-445, H_3_R-413, H_3_R-373, H_3_R-365, and H_3_R-329 cell lines. In contrast, much smaller FRET responses were observed for the H_3_R-415 and H_3_R-453, which might be related to lower expression levels of the latter (Supplementary Table 1).

### 3.3. Evaluation of ligand efficacies in inhibition of cAMP production by H_3_R isoforms

Next, all 18 reference H_3_R ligands were tested against all 7TM isoforms in the cAMP functional assay (Fig. 4). As expected, agonist-induced responses were highest for H_3_R-413 ≥ H_3_R-445 > H_3_R-329 > H_3_R-373 = H_3_R-365 = H_3_R-453 = H_3_R-415, whereas clear inverse agonism was observed for H_3_R-373 and H_3_R-365 isoforms (Figure 4s and 4t) followed by H_3_R-445 and H_3_R-329 isoforms (Fig. 4p and 4u). Interestingly, impentamine and dimethyl-impentamine acted as partial agonists on H_3_R-453, H_3_R-445, H_3_R-415, H_3_R-413, H_3_R-373, and H_3_R-329, but were full agonists on H_3_R-365 with the same intrinsic activity as histamine (Fig. 5).

**Fig. 4.**
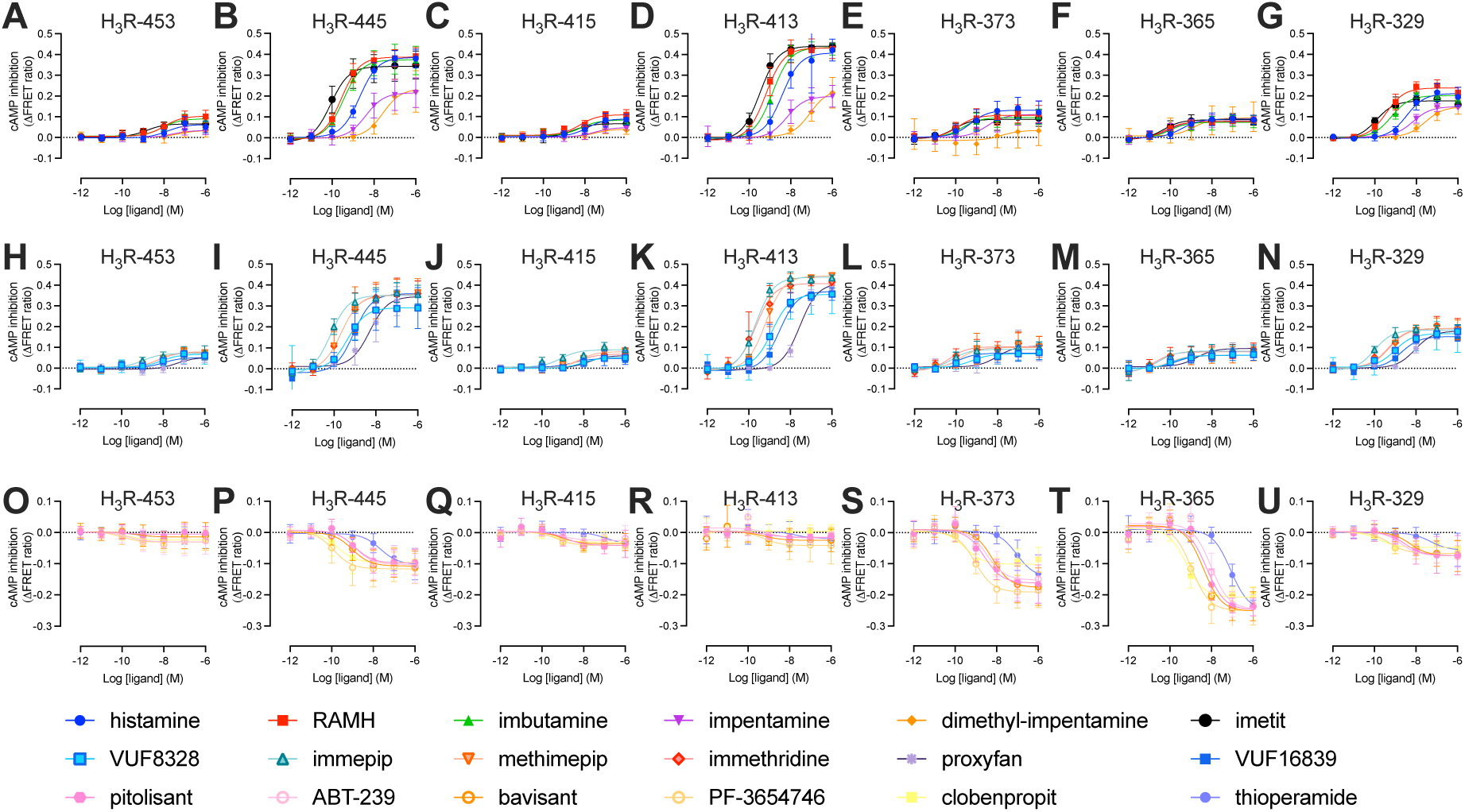
Concentration-response curves for the 18 reference ligands tested for their modulation of cAMP levels in HEK-EPAC cells stably expressing the seven H_3_R isoforms. The cells were pre-incubated with famotidine (1 µM) for 30 min and pre-stimulated with forskolin (10 µM) for 10 min. Seven concentrations of each ligand (agonists in panels a-n, inverse agonists in panels o-u) were added and cells were incubated for 90 min before the FRET signal of the EPAC sensor was measured. The data are mean ± S.D. from three or four independent experiments.

**Fig. 5.**
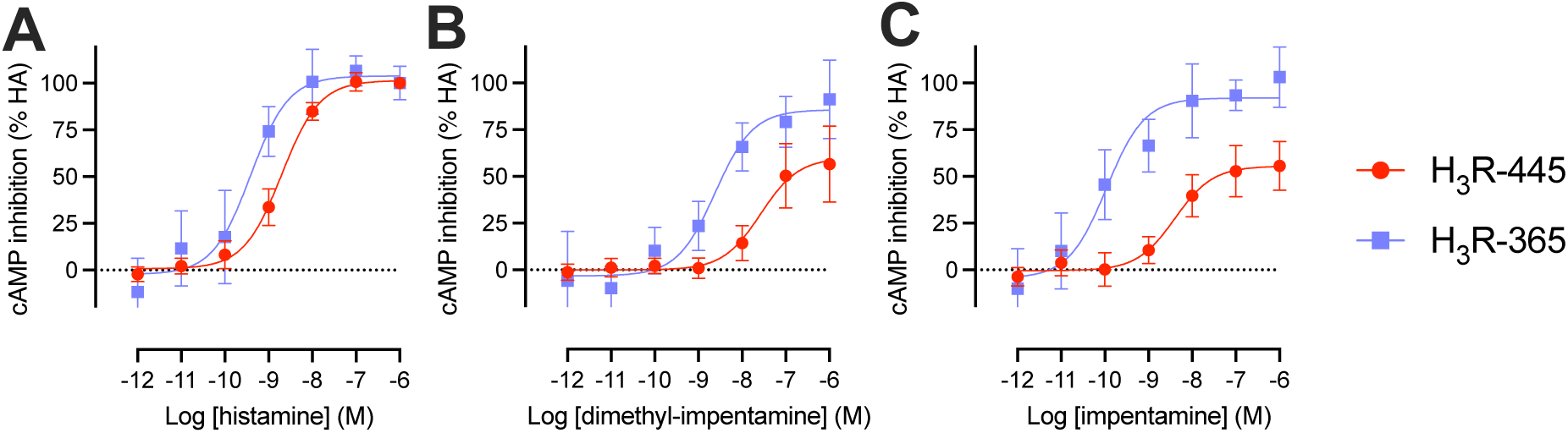
Comparison of the concentration response curves on H_3_R-445- and H_3_R-365-mediated inhibition of cAMP production. Concentration response curves of histamine (A), dimethyl-impentamine (B), and impentamine (C) on H_3_R-445 and H_3_R-365 (Figure 4B and 4F) are plotted as % histamine response for each isoform. The data are mean ± S.D. from three or four independent experiments performed in triplicate or duplicate.

The potency (pEC_50_) values observed for the H_3_R agonists in the EPAC biosensor assay generally correlated with their binding affinities for the seven H_3_R isoforms (Supplementary Fig. 4). However, different tendencies were observed between the H_3_R isoforms. All tested agonists showed lower pEC_50_ than pK_i_ values on H_3_R-453 and H_3_R-415, whereas this was also observed for most agonists on H_3_R-329 except for immepip, which has the same potency and affinity value for this shortest isoform. Conversely, apart from proxyfan, all agonists showed higher potency values for H_3_R-365-mediated signaling as compared to their affinities. Different agonist-specific signatures were observed on H_3_R-445, H_3_R-413, and H_3_R-373 with potencies (pEC_50_) being increased, unchanged, or decreased in comparison to the corresponding binding affinities (Fig. 6).

**Fig. 6.**
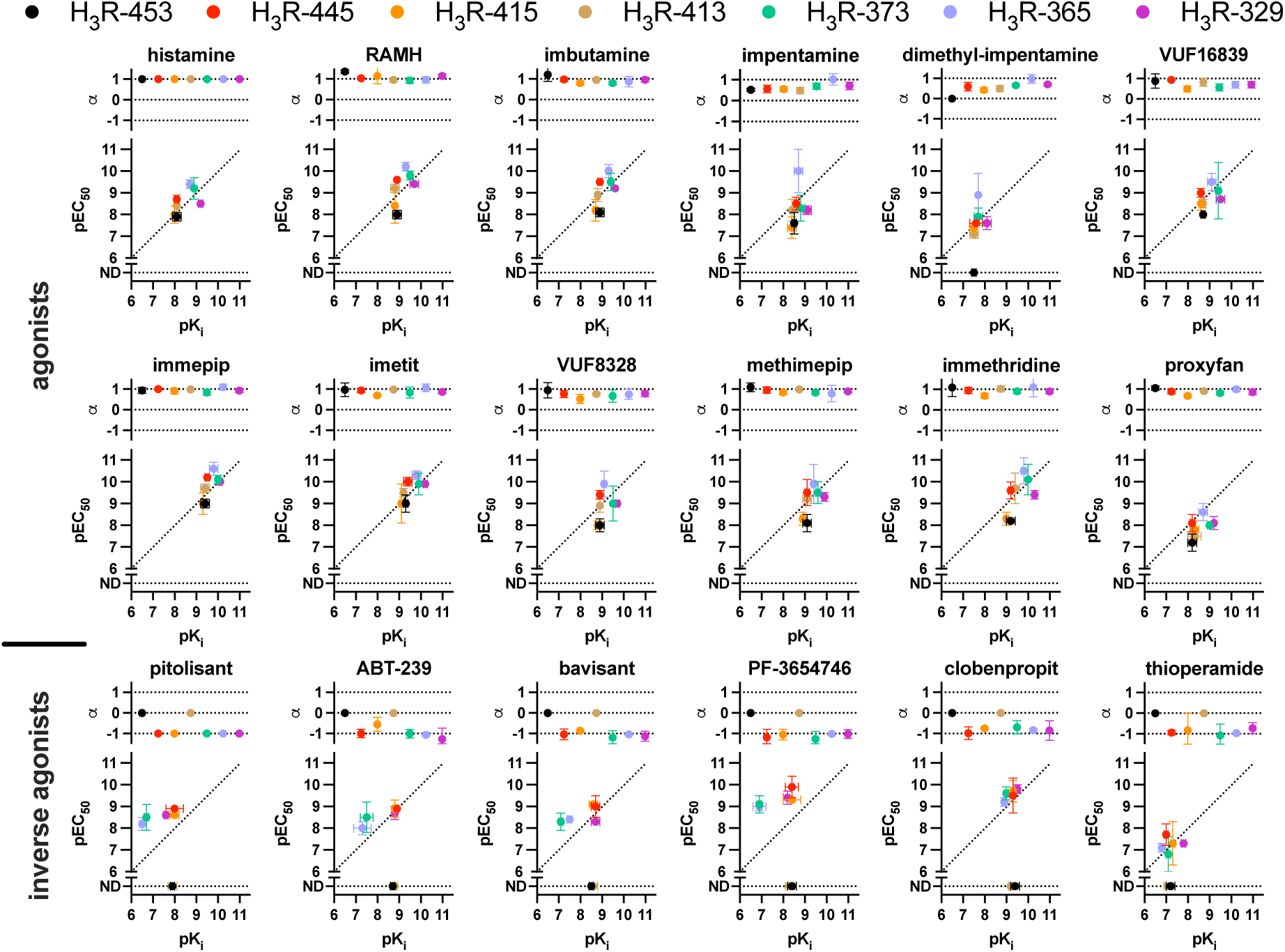
Summary of affinity, potency, and intrinsic activities of the 18 reference ligands on the seven H_3_R isoforms. The pK_i_ values derived from the competition binding experiments following Cheng-Prusoff conversion (Table 2) were plotted versus the potency (pEC_50_) values observed in the cAMP assay (Fig 4) for each ligand on the seven H_3_R isoforms, in combination with the corresponding intrinsic activity (α) values calculated relative to histamine and pitolisant for agonists and inverse agonists, respectively. Data are plotted as mean ± S.D. from at least three independent experiments. The line of unity is indicated with a dotted line and ND = not detectable.

**Fig. 7.**
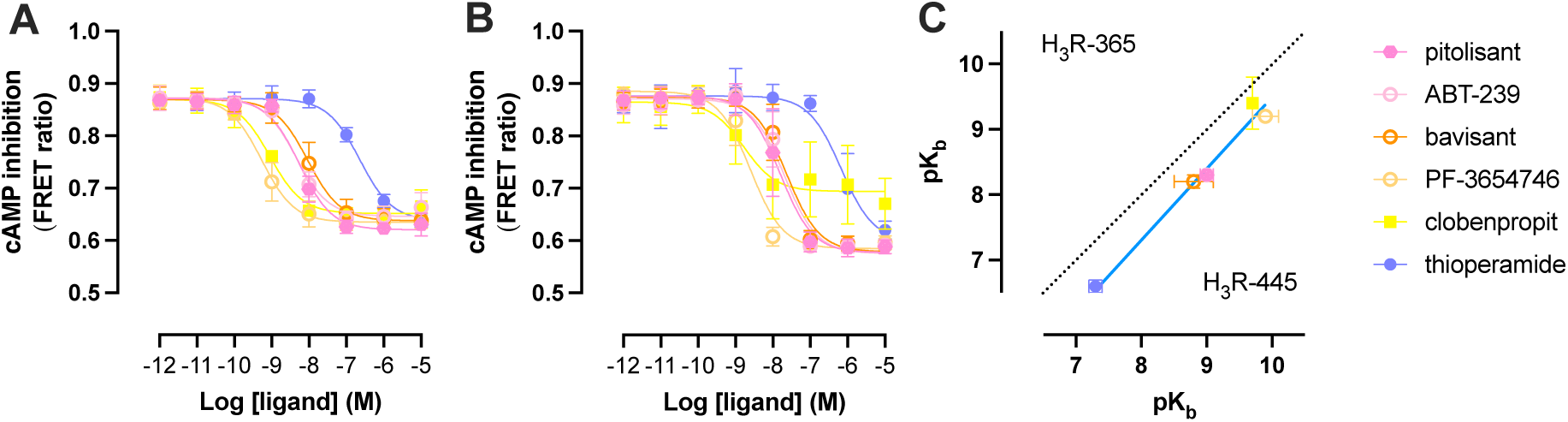
Antagonism of histamine-induced H_3_R-445- and H_3_R-365-mediated inhibition of cAMP production in HEK-EPAC cells. Cells were pre-incubated with famotidine (1 µM) for 30 min, pre-stimulated with forskolin (10 µM) for 10 min, and pre-incubated with increasing concentrations of inverse agonists for 30 min. Cells expressing H_3_R-445 (A) and H_3_R-365 (B) were stimulated with 8 and 1.6 nM histamine (i.e. EC_80_ concentrations), respectively, for 30 min before the FRET signal of the EPAC sensor was measured. The IC_50_ values of the inhibitory curves were converted into pK_b_ values using the Cheng-Prusoff correction and pK_b_ values for H_3_R-445 were plotted versus H_3_R-365 (C). Deming linear regression (blue line) was used to compare the pK_b_ values distribution to the (dotted) line of unity. The data are mean ± SD from three independent experiments.

### 3.4. Antagonism of histamine-induced H_3_R-445 and H_3_R-365 signaling

For the inverse agonists, the observed pEC_50_ values for the inhibition of constitutive receptor signaling were less well correlated to corresponding pK_i_ values for these different H_3_R isoforms (Supplementary Fig. 4). While pEC_50_ values of most inverse agonists were similar or only slightly increased compared to pK_i_ values for H_3_R-445, H_3_R-415, and H_3_R-329, much larger differences between potency and affinity were observed for pitolisant, ABT-239, bavisant, and PF-3654746 on the shorter and highly constitutively active H_3_R-365 and H_3_R-373 isoforms. In contrast, such an increase in potency values was not observed for clobenpropit and thioperamide in comparison to their affinity values for H_3_R-365 and H_3_R-373, which might be related to the much smaller difference in binding affinity between H_3_R-445 and both shorter isoforms for clobenpropit and thioperamide as compared to the other non-imidazole antagonists.

To evaluate the consequence of the affinity difference of the inverse agonists for canonical H_3_R-445 and the shorter H_3_R-365 isoform in the context of antagonizing histamine-induced receptor signaling, cells were stimulated with EC_80_ concentration histamine upon pre-incubation with increasing concentrations inverse agonist. The functional affinity (pK_b_) values confirm that the inverse agonists, except for clobenpropit, have significantly (unpaired t-test p<0.05) lower affinity for the more constitutively active short isoform H_3_R-365 and consequently higher concentrations (1.8- to 4.6-fold) are needed to inhibit histamine-induced receptor signaling.

## Discussion

Alternative splicing combines different exons or exon portions during RNA processing so that multiple protein isoforms can be encoded by a single gene in a cell-type specific manner (Wright et al., 2022). Approximately, half of the GPCR genes contain introns and are potentially subjected to alternative splicing, resulting in one or more isoforms with distinct pharmacological and/or signaling properties (Markovic and Challiss, 2009). A recent analysis of mRNA expression data from the Genotype-Tissue Expression (GTEx) consortium revealed that 42% of the 111 GPCRs with US Food and Drugs Administration (FDA)-approved drugs express more than one isoform with different tissue-specific (co-)expression profiles (Marti-Solano et al., 2020). Consequently, systemically administered drugs might induce tissue-specific responses by acting on differentially (co-)expressed isoforms (Marti-Solano et al., 2020). Interestingly, this study (https://gpcrdb.org/protein/isoforms; accessed 30 November 2023) reported only the reference H_3_R-445 and extended isoform H_3_R-453 (Marti-Solano et al., 2020), whereas five additional 7TM splice variants of the H_3_R have been previously reported that all share the same orthosteric ligand binding amino acid sequence with the canonical H_3_R-445 isoform but differ in the length of their ICL3 and/or intracellular C-terminal tail (Coge et al., 2001; Lovenberg et al., 1999; Nakamura et al., 2000; Tardivel-Lacombe et al., 2001; Wellendorph et al., 2002). Reverse transcription polymerase chain reaction analyses revealed that H_3_R isoforms are differentially expressed in various brain areas with generally higher mRNA levels for the H_3_R-445 and H_3_R-365 isoforms as compared to H_3_R-415, H_3_R-413, and H_3_R-329 (Coge et al., 2001; Esbenshade et al., 2006b; Wellendorph et al., 2002). Hence, these H_3_R isoforms could potentially be involved in the regulation of distinct neurological processes.

However, since the cloning of the H_3_R in the late 90s (Lovenberg et al., 1999), H_3_R drug discovery and lead optimization has been mostly focused on the canonical H_3_R-445 isoform. Hitherto, this resulted in a few dozen antagonists/inverse agonists being tested in clinical trial studies for various neurological disorders and the approval of pitolisant to treat narcolepsy (Wakix®) and more recently excessive daytime sleepiness in patients with obstructive sleep apnoea (Ozawade®)(Syed, 2016; Urquhart, 2019). Importantly, higher agonist binding affinities have been previously reported for the shorter H_3_R-365 isoform as compared to H_3_R-445, whereas an opposite change in binding affinities was observed for inverse agonists (Bongers et al., 2007b; Coge et al., 2001; Esbenshade et al., 2006a). Even though The H_3_R-365 isoform is more constitutive activity as compared to H_3_R-445 (Bongers et al., 2007a; Wellendorph et al., 2002), which corroborates with the paradigm that agonists and inverse agonists display higher binding affinities for active versus inactive receptor conformations, respectively (Staus et al., 2016). Indeed, opposite conformational changes have been detected with a bioluminescence resonance energy transfer (BRET)-based conformational H_3_R biosensor with HaloTag replacing ICL3 and the engineered luciferase NanoLuc fused to the C-tail upon stimulation with agonists and inverse agonists (Ma et al., 2022; Schihada et al., 2020). In the current study, tested agonists and antagonists/inverse agonists display similar binding affinities for the longer isoforms H_3_R-453, H_3_R-445, H_3_R-415, and H_3_R-413, which is in line with their comparable levels of basal receptor signaling and minor (or lack of) inverse agonist response (Fig. 3D and Fig. 4). Hence, the deletion of amino acid sequences ^234^D-Q^263^ or ^274^R-S^305^ in ICL3 of H_3_R-415 and H_3_R-413, respectively, or the 8-amino acid long ^446^KMKKKTCL^453^ C-tail extension in H_3_R-453 has no major effect on the conformational receptor state and consequently the ligand binding affinity for the orthosteric binding pocket.

As anticipated, all tested agonists have significantly higher affinities for the more constitutively active short isoforms H_3_R-365 and H_3_R-373, whereas lower binding affinities were observed for inverse agonists. The KMKKKTCL C-tail extension does not affect ligand binding affinities between both short isoforms, as also shown for H_3_R-445 and H_3_R-453. Surprisingly, agonists display increased binding affinities for the shortest H_3_R-329 isoform but its binding affinity for inverse agonists was found to be comparable to H_3_R-445. The 116-amino acid deletion in ICL3 of H_3_R-329 leaves only the C-terminal sequence SFTQ (i.e. amino acids 343-346 in canonical H_3_R-445 sequence) which might hamper this receptor isoform to adopt a conformation that supports high affinity inverse agonist binding, despite the fact that inverse agonism can be detected.

In contrast to the other five H_3_R isoforms, both H_3_R-365 and H_3_R-373 lack the SFTQRFRLSRD sequence (i.e. amino acids 343-353 in the canonical H_3_R-445 sequence) that constitutes the last 4 amino acids of ICL3 and two helical turns of the intracellular end of TM6 (Peng et al., 2022). The last amino acid of this deleted sequence in H_3_R-365 and H_3_R-373 corresponds to D^353^ at position 6.30x30 in the canonical H_3_R-445 sequence (https://gpcrdb.org/protein/hrh3_human/; accessed 30 November 2023). Importantly, aspartic or glutamic acid residues at position 6.30x30 have been identified to form an interhelical ionic lock with the highly conserved arginine at position 3.50x50 in the DRY motif in the intracellular end of TM3 to stabilize inactive receptor conformations (Palczewski et al., 2000). Site-directed mutagenesis or genetic variants of aspartic or glutamic acid at 6.30x30 have been found to disrupt this ionic lock in many different GPCRs such as for example histamine H_1_ receptor (Ma et al., 2021), β_2_-adrenergic receptor (Ballesteros et al., 2001), serotonin 5HT_2A_ receptor (Shapiro et al., 2002), resulting in increased constitutively signaling and associated changes ligand binding affinities. Hence, the absence of an ionic lock between TM3 and TM6 is hypothesized to cause the higher constitutive activity of H_3_R-365 and H_3_R-373.

In addition to the reference isoform H_3_R-445, only the H_3_R-413 isoform has also been detected in mouse, rat, guinea pig, and monkey (Ding et al., 2005; Drutel et al., 2001; Lovenberg et al., 2000; Morisset et al., 2000; Morisset et al., 2001; Rouleau et al., 2004; Strakhova et al., 2008; Tardivel-Lacombe et al., 2000; Yao et al., 2003). Importantly, the rat H_3_R-413 isoform was proposed to act as autoreceptor on histaminergic neurons based on a difference in stereoselectivity for the chiral ligands N^α^-methyl-α-chloromethylhistamine and sopramidine as compared to the rat H_3_R-445, whereas the latter was proposed to function as post-synaptic or heteroreceptor on other neurons (Gbahou et al., 2012). The extended and shorter human isoforms H_3_R-453, H_3_R-415, H_3_R-373, H_3_R-365, and H_3_R-329 have so far not been identified in these species, which might be related to lower transcript levels as compared to longer isoforms preventing their readily detection by reverse transcription polymerase chain reaction (Ding et al., 2005; Yao et al., 2003). On the other hand, various additional isoforms have been found in mouse (H_3_R-397), rat (H_3_R-410 and H_3_R-397), monkey (H_3_R-410) and hamster (H_3_R-406) that have not been detected in human (Barrett et al., 2005; Morisset et al., 2001; Rouleau et al., 2004; Strakhova et al., 2008). Interestingly, the hamster H_3_R-406 has a similar C-terminal tail extension (KMEEKKTRL) as observed in human H_3_R-445 and H_3_R-373. Considering that the three shorter human isoforms showed displayed largest differences in ligand binding and/or constitutive activity as compared to reference isoform H_3_R-445, it would be interesting to investigate whether these shorter isoforms do exist in these species that are used for disease models in pre-clinical drug testing and where they are localized in the central nervous system. In contrast to human H_3_R-445, the shorter H_3_R-365 is not able to stimulate glycogen synthase kinase 3β (GSK3β) phosphorylation, while also difference signaling bias profiles were observed for 5 agonists at these two isoforms (Riddy et al., 2017), whereas analysis of a larger set of agonists on H_3_R-445, H_3_R-415, H_3_R-365, and H_3_R-329 revealed distinct agonist efficacies in G_i_-mediated signaling between isoforms (Rahman et al., 2023).

In conclusion, for future H_3_R drug discovery, more detailed analysis of the spatial expression and signaling capacities of all H_3_R isoforms is needed as well as the effect of inverse agonists on basal and histamine-induced signaling by H_3_R isoforms to relevant functional responses considering their opposite change in binding affinities on the more constitutive active shorter isoforms H_3_R-365 and H_3_R-373. Moreover, such analysis might identify potential signaling biased ligands as previously reported for the β-arrestin2-biased reference antagonists JNJ7777120 on the histamine H_4_ receptor (Nijmeijer et al., 2012; Nijmeijer et al., 2013; Rosethorne et al., 2016).

## CRediT authorship contribution statement

**Meichun Gao**: investigation, methodology, writing – original draft, formal analysis, validation, funding acquisition. **Mabel Dekker**: investigation, methodology, writing – original draft, formal analysis. **Rob Leurs**: conceptualization, supervision, methodology, writing – review & editing, project administration, resources. **Henry Vischer**: conceptualization, supervision, methodology, writing – review & editing, project administration, resources.

## Acknowledgements

Meichun Gao is supported by CSC Chinese scholarship grant [202006310027]. We thank Dr. Mirjam Zimmermann for providing the HEK293-EPAC cells.

## Notes

### Competing Interest Statement

The authors have declared no competing interest.

