## supplemental data for "Pharmacological characterization of seven human histamine H_3_ receptor isoforms"

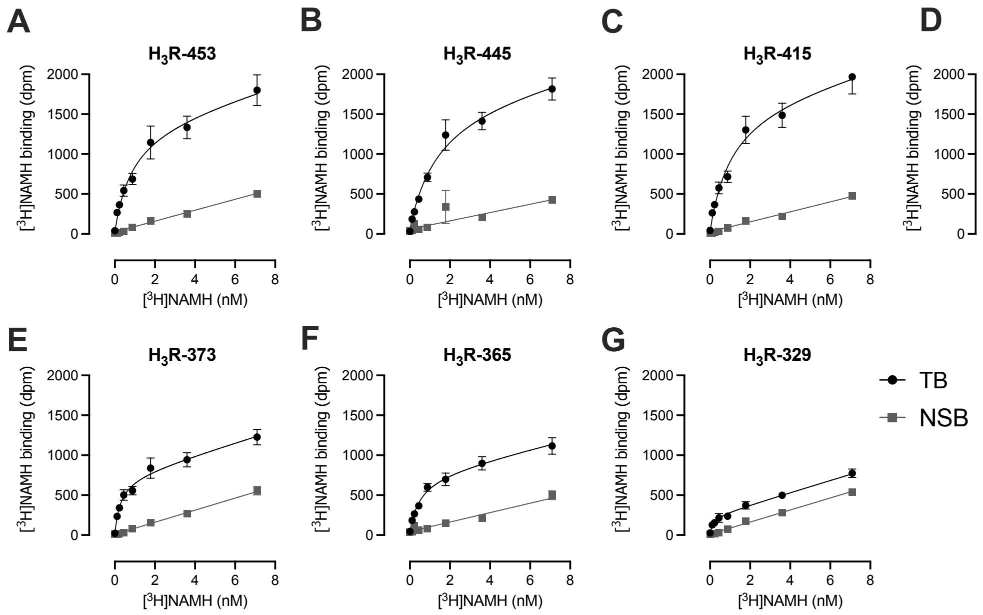


**Supplementary Fig. 1.** Saturation binding of radioligand agonist [$^3$H]NAMH to $H_3$R isoforms. Total (TB) and non-specific (NSB) binding of [$^3$H]NAMH to HEK293T membranes transiently expressing $H_3$R-453 (A), $H_3$R-445 (B), $H_3$R-415 (C), $H_3$R-413 (D), $H_3$R-373 (E), $H_3$R-365 (F), and $H_3$R-329 (G), was measured in the absence or presence of 10 µM clobenpropit, respectively. Data are show as mean ± SEM of 3 or 4 independent experiments performed in triplicate.

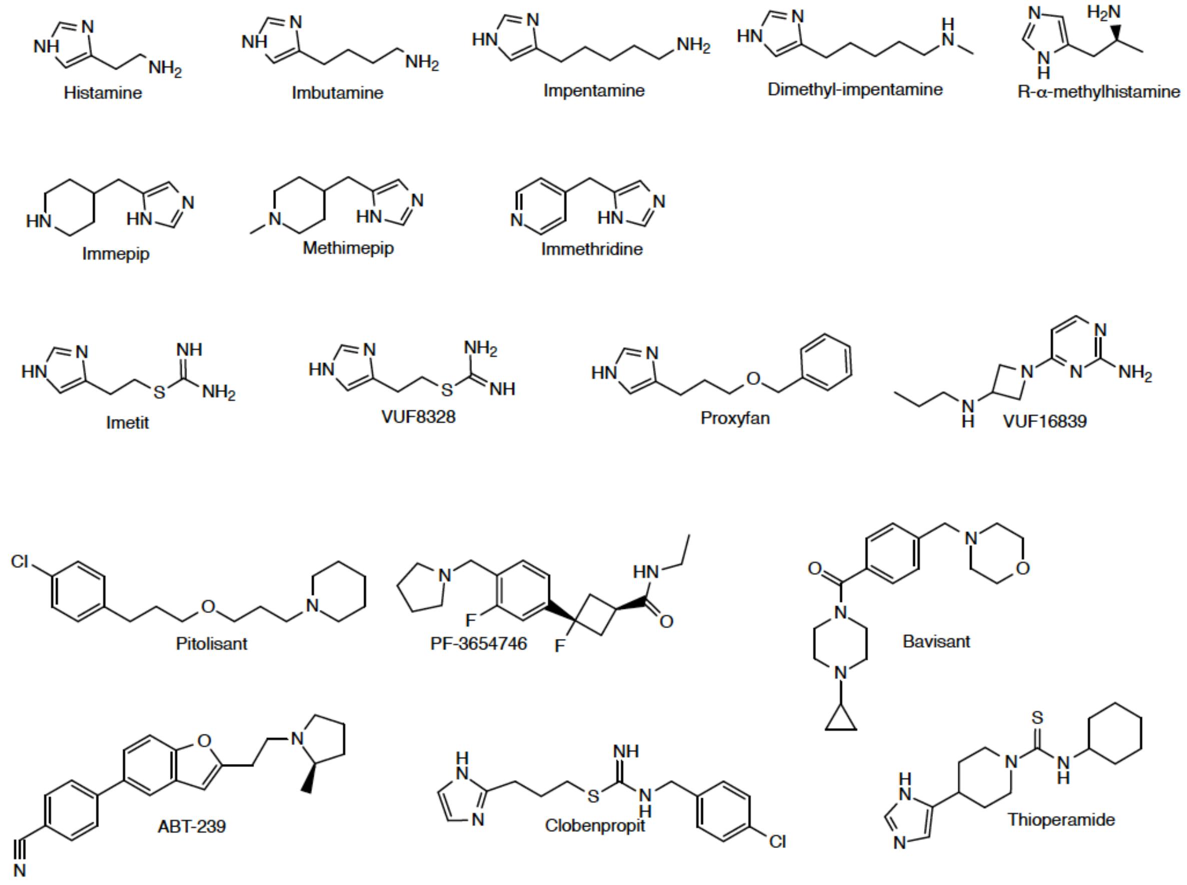


**Supplementary Fig. 2.** Structures of 18 reference $H_3R$ agonists and antagonists/inverse agonists that were evaluated in this study.

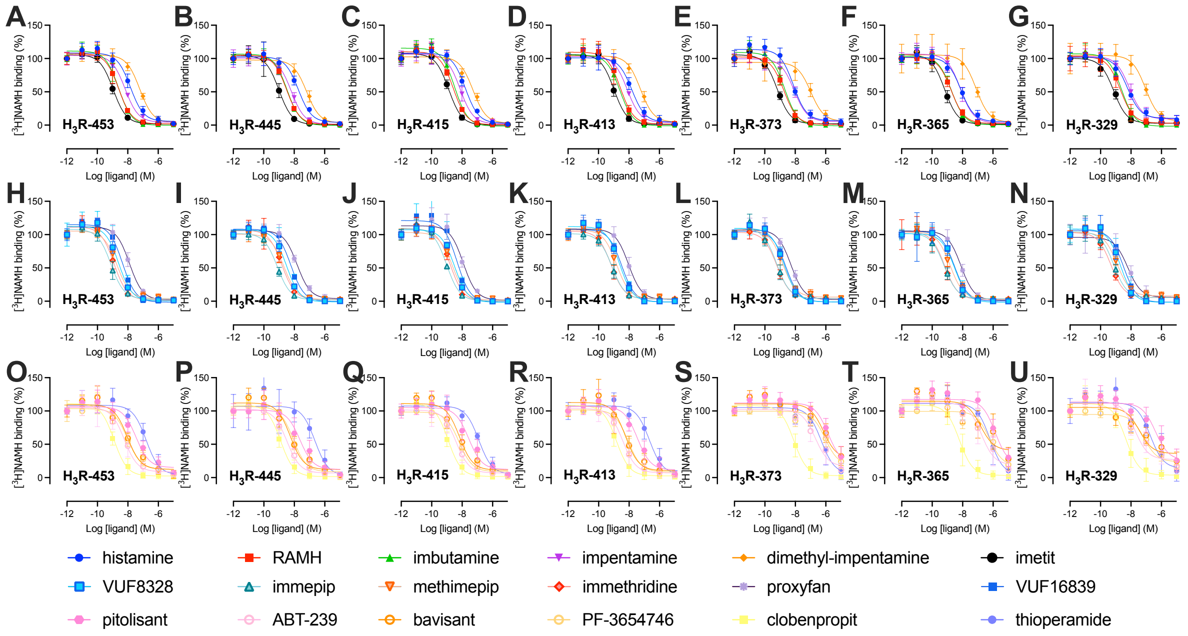


**Supplementary Fig. 3. Competition binding between radioligand [$^{3}$H]NAMH and 18 reference $H_3R$ ligands to the seven $H_3R$ isoforms.** Binding of 2 nM [3H-NAMH] to HEK293T membranes transiently expressing $H_3R$-453 (A, H, O), $H_3R$-445 (B, I, P), $H_3R$-415 (C, J, Q), $H_3R$-413 (D, K, R), $H_3R$-373 (E, L, S), $H_3R$-365 (F, M, T), and $H_3R$-329 (G, N, U), was measured in the presence of increasing concentration unlabeled $H_3R$ agonist (A-N) or antagonists/inverse agonists (O-U). Data are show as mean ± SD of at least 3 independent experiments performed in triplicate.

**Supplementary Table 1.** $H_3R$ Isoform expression as measured by [$^{3}$H]NAMH binding in monoclonal HEK-EPAC cell lines.

| EPAC cell line | $B_{max}$ level (fmol/mg protein) |
|---|---|
| $H_3R$-453 | 148 ± 64 |
| $H_3R$-445 | 561 ± 213 |
| $H_3R$-415 | 336 ±110 |
| $H_3R$-413 | 719 ± 30 |
| $H_3R$-373 | 475 ± 137 |
| $H_3R$-365 | 1024 ± 14 |
| $H_3R$-329 | 427 ± 42 |

Data shown are mean ± SD of 3 independent experiments. Brown-Forsythe and Welch ANOVA followed by a Dunnett's multiple T3 comparisons test showed no statistical difference ($p > 0.05$) in Bmax values between $H_3R$-445 and the other isoforms.

**Supplementary Table 2. The potency ($pEC_{50}$) and intrinsic activity (α) values of 18 ligands as measured in the EPAC biosensor assay for in HEK293 cells stably transfected with $H_3R$-453, -445, -415, -413, -373, -365 or -329 isoforms.**

| | $H_3R$-453 | | $H_3R$-445 | | $H_3R$-415 | | $H_3R$-413 | | $H_3R$-373 | | $H_3R$-365 | | $H_3R$-329 | |
|---|---|---|---|---|---|---|---|---|---|---|---|---|---|---|
| | $pEC_{50}$ | α | $pEC_{50}$ | α | $pEC_{50}$ | α | $pEC_{50}$ | α | $pEC_{50}$ | α | $pEC_{50}$ | α | $pEC_{50}$ | α |
| **Histamine** | 7.9 ± 0.2* | 1.0 ± 0.0 | 8.7 ± 0.1 | 1.0 ± 0.0 | 8.0 ± 0.4 | 1.0 ± 0.0 | 8.4 ± 0.5 | 1.0 ± 0.0 | 9.2 ± 0.5 | 1.0 ± 0.0 | 9.4 ± 0.2 | 1.0 ± 0.0 | 8.5 ± 0.1 | 1.0 ± 0.0 |
| **Immepip** | 9.0 ± 0.2* | 0.9 ± 0.1 | 10.2 ± 0.1 | 1.0 ± 0.0 | 9.0 ± 0.5 | 0.9 ± 0.1 | 9.7 ± 0.2 | 1.0 ± 0.0 | 10.1 ± 0.2 | 0.8 ± 0.2 | 10.6 ± 0.2 | 1.1 ± 0.1 | 10.0 ± 0.1 | 0.9 ± 0.1 |
| **R-α-methyl histamine (RAMH)** | 8.0 ± 0.2* | 1.4 ± 0.1 | 9.6 ± 0.1 | 1.0 ± 0.0 | 8.4 ± 0.8 | 1.1 ± 0.4 | 9.2 ± 0.2 | 1.0 ± 0.1 | 9.8 ± 0.2 | 0.9 ± 0.1 | 10.2 ± 0.2 | 1.0 ± 0.2 | 9.4 ± 0,1 | 1.1 ± 0.1 |
| **Imbutamine** | 8.1 ± 0.2* | 1.2 ± 0.3 | 9.5 ± 0.1 | 1.0 ± 0.1 | 8.2 ± 0.5 | 0.8 ± 0.1 | 8.9 ± 0.3 | 0.9 ± 0.0 | 9.5 ± 0.4 | 0.8 ± 0.1 | 10.0 ± 0.3 | 0.9 ± 0.3 | 9.2 ± 0.1 | 1.0 ± 0.1 |
| **Impentamine** | 7.6 ± 0.5 | 0.5 ± 0.1 | 8.5 ± 0.3 | 0.6 ± 0.2 | 7.4 ± 0.5 | 0.5 ± 0.1 | 8.2 ± 0.5 | 0.5 ± 0.1 | 8.3 ± 0.6 | 0.7 ± 0.1 | 10.0 ± 1.0 | 1.0 ± 0.3 | 8.2 ± 0.2 | 0.7 ± 0.2 |
| **dimethyl-impentamine** | NE | NE | 7.6 ± 0.1 | 0.6 ± 0.2 | 7.4 ± 0.2 | 0.4 ± 0.1 | 7.1 ± 0.2* | 0.5 ± 0.2 | 7.9 ± 0.4 | 0.7 ± 0.1 | 8.9 ± 1.0 | 1.0 ± 0.2 | 7.6 ± 0.3 | 0.7 ± 0.1 |
| **Imetit** | 9.0 ± 0.4 | 1.0 ± 0.3 | 10.0 ± 0.2 | 0.9 ± 0.1 | 9.0 ± 0.9 | 0.7 ± 0.0 | 9.5 ± 0.2 | 1.0 ± 0.0 | 9.9 ± 0.5 | 0.8 ± 0.3 | 10.3 ± 0.2 | 1.1 ± 0.2 | 9.9 ± 0.0 | 0.9 ± 0.1 |
| **VUF8328** | 8.0 ± 0.3* | 0.9 ± 0.4 | 9.4 ± 0.2 | 0.8 ± 0.2 | 8.0 ± 0.2* | 0.5 ± 0.2 | 8.9 ± 0.3 | 0.8 ± 0.1 | 9.0 ± 0.8 | 0.7 ± 0.3 | 9.9 ± 0.6 | 0.7 ± 0.2 | 9.0 ± 0.0 | 0.8 ± 0.2 |
| **Methimepip** | 8.1 ± 0.4 | 1.1 ± 0.2 | 9.5 ± 0.6 | 1.0 ± 0.2 | 8.3 ± 0.3 | 0.8 ± 0.1 | 9.2 ± 0.3 | 1.0 ± 0.1 | 9.5 ± 0.5 | 0.8 ± 0.1 | 9.9 ± 0.9 | 0.8 ± 0.4 | 9.3 ± 0.2 | 0.9 ± 0.1 |
| **Immethridine** | 8.2 ± 0.1* | 1.1 ± 0.4 | 9.6 ± 0.4 | 0.9 ± 0.2 | 8.3 ± 0.3* | 0.7 ± 0.1 | 9.7 ± 0.7 | 1.0 ± 0.1 | 10.1 ± 0.7 | 0.9 ± 0.1 | 10.5 ± 0.5 | 1.1 ± 0.5 | 9.4 ± 0.2 | 0.9 ± 0.1 |
| **Proxyfan** | 7.2 ± 0.4 | 1.0 ± 0.2 | 8.1 ± 0.4 | 0.9 ± 0.1 | 7.8 ± 0.4 | 0.7 ± 0.1 | 7.5 ± 0.2 | 0.9 ± 0.1 | 8.0 ± 0.0 | 0.8 ± 0.1 | 8.6 ± 0.3 | 1.0 ± 0.1 | 8.1 ± 0.3 | 0.9 ± 0.1 |
| **VUF16839** | 8.0 ± 0.1* | 0.9 ± 0.3 | 9.0 ± 0.2 | 0.9 ± 0.0 | 8.5 ± 0.3 | 0.5 ± 0.1 | 8.5 ± 0.2 | 0.8 ± 0.2 | 9.1 ± 1.3 | 0.6 ± 0.2 | 9.5 ± 0.4 | 0.7 ± 0.2 | 8.7 ± 0.1 | 0.7 ± 0.2 |
| **Pitolisant** | NE | NE | 8.9 ± 0.1 | -1.0 ± 0.0 | 8.6 ± 0.1* | -1.0 ± 0.0 | NE | NE | 8.5 ± 0.6 | -1.0 ± 0.0 | 8.2 ± 0.3 | -1.0 ± 0.0 | 8.6 ± 0.1 | -1.0 ± 0.0 |
| **ABT-239** | NE | NE | 8.9 ± 0.1 | -1.0 ± 0.2 | 8.9 ± 0.4 | -0.5 ± 0.3 | NE | NE | 8.5 ± 0.7 | -1.0 ± 0.2 | 8.0 ± 0.3 | -1.1 ± 0.0 | 8.7 ± 0.3 | -1.3 ± 0.5 |
| **Bavisant** | NE | NE | 9.0 ± 0.5 | -1.0 ± 0,3 | 9.1 ± 0.1 | -0.9 ± 0.0 | NE | NE | 8.3 ± 0.4 | -1.2 ± 0.3 | 8.4 ± 0.1 | -1.0 ± 0.0 | 8.3 ± 0.1 | -1.1 ± 0.2 |
| **PF-3654746** | NE | NE | 9.9 ± 0.5 | -1.2 ± 0.4 | 9.3 ± 0.1 | -1.1 ± 0.3 | NE | NE | 9.1 ± 0.4 | -1.3 ± 0.4 | 9.0 ± 0.1 | -1.0 ± 0.1 | 9.4 ± 0.3 | -1.0 ± 0.2 |
| **Clobenpropit** | NE | NE | 9.5 ± 0.8 | -1.0 ± 0.3 | 9.7 ± 0.5 | -0.8 ± 0.0 | NE | NE | 9.6 ± 0.3 | -0.7 ± 0.3 | 9.2 ± 0.2 | -0.8 ± 0.1 | 9.8 ± 0.2 | -0.9 ± 0.5 |
| **Thioperamide** | NE | NE | 7.7 ± 0.5 | -0.9 ± 0.1 | 7.3 ± 1.0 | -0.8 ± 0.8 | NE | NE | 6.8 ± 0.8 | -1.1 ± 0.6 | 7.1 ± 0.2 | -1.0 ± 0.0 | 7.3 ± 0.1 | -0.7 ± 0.3 |

For each isoform, histamine was used as full agonist and pitolisant as full inverse agonist to calculate the intrinsic activities(α) of the other ligands at that specific isoform. The data are mean ± SD from three or four independent experiments. Statistical difference ($p<0.05$) in $pEC_{50}$ values for the various isoforms in comparison to $H_3R$-445 was determined using Brown-Forsythe and Welch ANOVA followed by a Dunnett’s multiple T3 comparisons test and is indicated with an asterisk (*). NE = no effect.

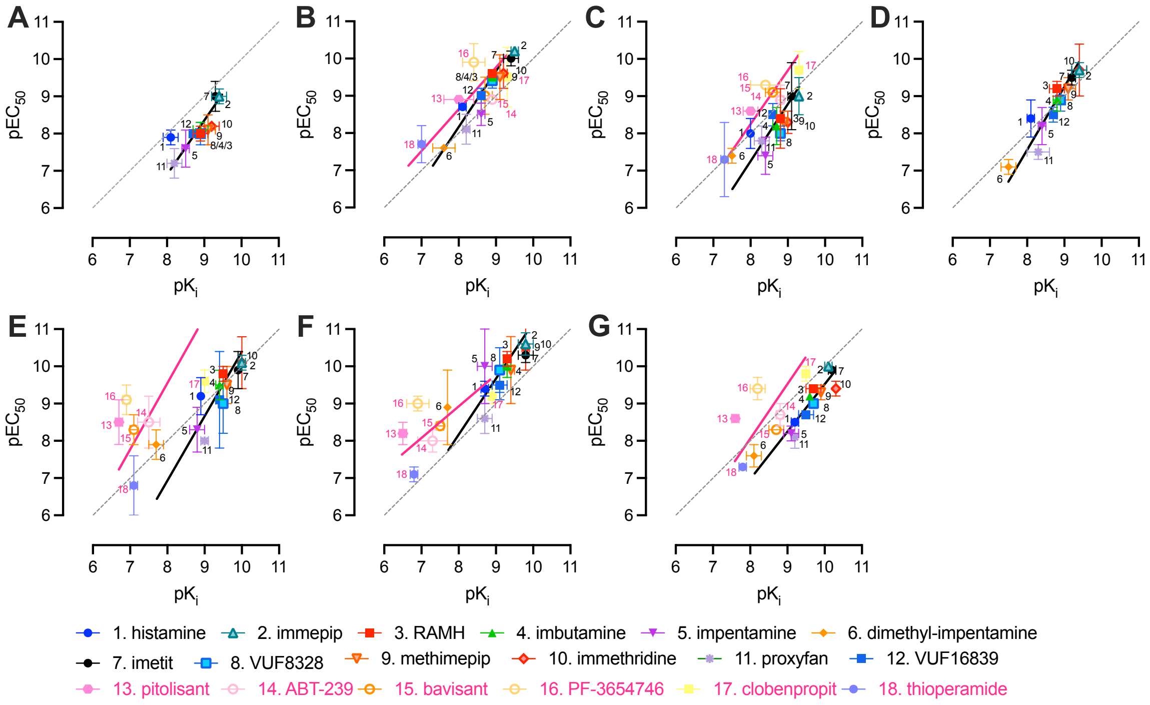


**Supplementary Fig. 4. Comparison ligand affinity and potency values on $H_3R$ isoforms.** The $pK_i$ values obtained by radioligand competition binding assays for $H_3R$-453 (A), $H_3R$-445 (B), $H_3R$-415 (C), $H_3R$-413 (D), $H_3R$-373 (E), $H_3R$-365 (F), and $H_3R$-329 (G) are plotted versus and the $pEC_{50}$ values obtained with the EPAC biosensor assay. Agonist and inverse agonist are numbered in black and magenta, respectively. Data are mean of three independent experiments performed in triplicate or duplicate. Deming linear regression was used to compare the affinity and potency, the dotted line represents line of unity.
